# Poor protection of amphibian evolutionary history reveals opportunities for global protected areas

**DOI:** 10.1101/2020.10.15.338061

**Authors:** Jasmin Upton, Claudia L. Gray, Benjamin Tapley, Kris A. Murray, Rikki Gumbs

## Abstract

As habitat loss is a major driver of amphibian population declines, protected areas (PAs) can play a crucial role in amphibian conservation. Documenting how well the global PA network captures the evolutionary history of amphibians can inform conservation prioritisation and action. We conducted a phylogenetic gap analysis to assess the extent to which amphibian phylogenetic diversity (PD) is unprotected by the PA network and compared this to other terrestrial vertebrate groups. 78% of amphibian species and 64% of global amphibian PD remains unprotected, which is higher than corresponding figures for squamates, mammals and birds. Amongst amphibians, salamanders were the least well protected, with 78% of PD unprotected, compared with 64% for caecilians and 63% for frogs. We identify areas that offer the greatest opportunity to capture unprotected amphibian evolutionary history. We could capture an additional 29.4% of amphibian PD, representing 40 billion years of evolutionary history, by protecting an additional 1.9% of global amphibian distributions (1.74% of global land area) and increasing the restrictions in 0.6% of amphibian distributions to match the management objectives of PAs in IUCN categories I or II. Importantly, we found that the spatial distribution of unprotected PD was correlated across all groups, indicating that expanding the PA network to conserve amphibian PD can secure imperilled vertebrate diversity more generally.

## 1. Introduction

Amphibian declines are a global conservation crisis. Approximately 41% of assessed amphibians are threatened (excluding Data Deficient assessments), compared with 14% of birds, 19% of reptiles and 26% of mammals (Böhm et al. 2013, IUCN 2020). The Global Amphibian Assessment found 43% of amphibian species to be under rapid demographic decline, and 7.4% of species facing imminent extinction (Stuart et al. 2004). Global amphibian declines are driven by numerous, often synergistic, threats including habitat loss, infectious disease, invasive species and overexploitation (Stuart et al. 2004). The loss of amphibians can drastically affect food chains (Zipkin et al. 2020), alter nutrient exchange between aquatic and terrestrial systems and lower ecosystem biomass (Blaustein 1994; Colón-Gaud et al. 2009). In addition, there are strong biases in protection across taxa, tending to favour birds and mammals, and large gaps exist in our knowledge of the extinction risk faced by amphibians (Böhm et al. 2013; Meiri & Chapple 2016; Tapley et al. 2018).

Protected areas (PAs) can be an effective step in safeguarding biodiversity (Gray et al. 2016; Pacifici et al. 2020). Effective PAs could prove critical in amphibian conservation in the long term, as habitat loss poses the greatest threat to amphibians worldwide. Most amphibians are poor dispersers and range restricted and are therefore particularly sensitive to anthropogenic impacts on their habitats (Gardner et al. 2007; Chanson et al. 2008). However, the efficacy of PAs in protecting biodiversity is limited by their placement in areas that are not of high conservation value (Visconti et al. 2019). The successors to the Aichi targets, due to be agreed by the Convention on Biological Diversity in 2021, have the potential to motivate improvements in the distribution and extent of the PA network if based on scientific information.

The conservation of phylogenetic diversity (PD) is increasing in value within the global agenda. The Intergovernmental Science-Policy Platform on Biodiversity and Ecosystem Services (IPBES) recognises PD as a key indicator of nature’s contributions to people (IPBES 2019). Research highlighting gaps and opportunities for conserving PD is therefore currently of great importance. Faith’s (1992) metric of PD provides a measure to approximate feature diversity (Faith 1992; Forest et al. 2007) by summing the phylogenetic branch lengths connecting all species in a clade or set of taxa across a phylogenetic tree. Using this information, we can identify clades which have a disproportionately large contribution to global evolutionary history (Faith 1992).

The combination of spatial patterns of PD with measures of extinction risk or protection can guide prioritisations for the conservation of global diversity (Rosauer et al. 2017; Pollock et al. 2017). Gap analyses can be used to identify areas that contain disproportionately high amounts of unprotected PD not captured by the PA network (Scott et al. 1994; Rodrigues et al. 2004a); these are obvious priorities if PA expansion is to safeguard unique evolutionary history.

Here, we use gap analysis methods to identify branches of the terrestrial vertebrate tree of life that are not captured by the current terrestrial PA network. We provide a global assessment of unprotected amphibian PD and explore the differences between the three amphibian orders. We contrast results for amphibians with those for birds, mammals and squamates to determine whether amphibians are equitably protected by the current PA network. Finally, we identify grid cells where increased protection would provide the greatest potential gains for conserving global amphibian PD and demonstrate that large gains in safeguarding unique evolutionary history can be achieved with relatively small PA increases.

## 2. Methods

### 2.1. Species data

Spatial data were taken from IUCN (2017) for 6457 amphibian species (~80% of species, Frost 2020) and 5371 terrestrial mammal species (~83%, IUCN 2020), from BirdLife International (2017) for 9761 bird species (~88%, IUCN 2020) and from Roll et al. (2017) for 9557 squamates (~92%, Uetz & Hosek 2018). Amphibian orders were also assessed individually with spatial data downloaded from IUCN (2018). Distribution data were available for 5807 Anurans (frogs and toads; ~81%, Frost 2020), 562 Caudata (salamanders and newts; ~76%, Frost 2020) and 161 Gymnophiona (caecilians; 75%, Frost 2020). Only extant, resident ranges were used for amphibians, mammals and squamates and only extant, resident and breeding ranges were used for birds. Invasive ranges, where designated, were excluded and all distribution data were restricted to land.

We rasterized all distribution data at two spatial resolutions of ~1° (100 × 100 km) and ~2° (200 × 200 km) grid cells using a Mollweide equal area projection. Using too fine a spatial resolution can be misleading and increase the error in the correct placing of species ranges (Hurlbert & Jetz 2007); however, using too coarse a spatial resolution can misinterpret the distributions of very small ranging species and lead to spatial smoothing of the data and overestimated Extent Of Occurrence (EOO; Dormann et al. 2007). Differences in results under the two different spatial resolutions were negligible, so we present results at ~1° resolution in the main text and at ~2° resolution in the supporting information (**Figure A1 & A2**).

Phylogenetic data were taken from Jetz & Pyron (2018) for amphibians, Jetz et al. (2014) for birds, Kuhn et al. (2011) for mammals and Tonini et al. (2016) for squamates. Only taxa that were represented in the phylogeny and distribution data were included in the analyses, or 4374 mammals (67% of species), 7177 birds (64%), 9229 for squamates (89%), 5835 amphibians (72%). Within amphibians: 150 Gymnophiona (70%), 525 Caudata (71%), 5188 Anura (72%). Phylogenetic uncertainty in the available data was accounted for by randomly sampling 25 phylogenetic trees for each taxonomic group from the published ‘pseudoposterior’ distributions (Thomas et al. 2013), which are considered equally probable estimations of the phylogenetic relationships between species and clades. A sample of 25 trees was considered sufficient as there was little variation in the amount of PD unprotected on a global scale across all trees (**Figure A3**).

### 2.2. Protected area data

Spatial data for the global PA network were downloaded from the World Database on Protected Areas (IUCN & UNEP-WCMC 2019). Only PAs with polygon data that had a reported area larger than 0 km^2^ and a recognised terrestrial status (including designated, established and inscribed) were used. PAs with point data only were excluded from all analyses. Jones et al. (2018) found that more strictly managed PAs were subject to significantly lower levels of human pressure than the remaining categories, thus we first ran the analyses using only PAs in IUCN management category I and II, giving us a conservative estimate of the proportion of PD under protection. To evaluate the effect of our strict inclusion criteria for PAs, all analyses were repeated using all categories of IUCN PAs (Management categories I-VI) as well as all PAs labelled as ‘Not Applicable’, ‘Not Assigned’ and ‘Not reported’ to examine the increase in protected PD with less stringent protection criteria.

As 64.7% (12 714 of 19 637) of all PAs in IUCN management category I and II had a reported area less than or equal to 10 km^2^, a fine-scale resolution was essential for mapping the PA data, to reduce error in the detection of smaller PAs. All PAs were therefore rasterized at a resolution of 2.5 × 2.5 km and aggregated and reprojected with bilinear interpolation to match the resolution and extent of the amphibian range data. PAs outside the extent of amphibian distributions were excluded from the analyses.

We used a binary approach to determine whether a grid cell in the species range data was protected or unprotected (i.e. we did not consider any variation in the effectiveness of protection in each “protected” cell). We overlapped the fine-resolution PA raster with the lower resolution species raster and determined the proportion of each larger grid cell that was covered by PAs at the 2.5 × 2.5 km resolution (McGowan et al. 2020). We determined whether the grid cell was protected or not using eleven protection thresholds (PTs) ranging between 0 and 100%, at 10% intervals. The percentage of overlap between a larger grid cell and the PA polygon must meet or exceed the given PT to be considered protected. The 20% threshold is met when at least 20% of the larger grid cell overlaps with 2.5 × 2.5km PA grid cells (hereafter >20%). A broad interval range was chosen to reflect all possible scenarios of protection, ranging from an optimistic scenario (>0% threshold) to a more conservative scenario (100% overlap threshold).

We considered a species present in a grid cell if any of its range overlapped with the grid cell (Safi et al. 2013; Roll et al. 2017) and we considered a species protected if its range was found to occur in at least one protected grid cell (Rodrigues et al. 2004b). An alternative method would have been to scale protection based on the range size of the species, e.g. set a more demanding representation target (a larger percentage of the range) for species with more restricted ranges (Rodrigues et al. 2004a; Thuiller et al. 2015; Rosauer et al. 2017). However, we considered one grid cell of a species range as protected to be sufficient for the protection of the species, given that amphibians typically occur in just 1-2 grid cells (González-del-Pliego et al. 2019). The total number of protected and unprotected species for each taxonomic group was calculated. Results are based on the assumption that PAs are protecting species and, therefore, PD.

Two forms of error may occur from using a binary approach: 1) species considered absent from a PA are actually present within it (false absence), i.e. where the proportion of overlap of the PA with the grid cell does not meet the PT so the grid cell is considered unprotected; and 2) where species considered present within a PA are actually absent (false presence), i.e. the range of the species within a protected grid cell, does not actually overlap with the range of the PA (Rodrigues et al. 2004a). Selecting for larger PTs decreases the likelihood of both errors occurring; however, higher PTs leave fewer grid cells protected and reduce our ability to observe the effects of PAs (**Figure A4**). To balance these sources of error, and as a somewhat arbitrary selection after observing the accumulation curves of PD protection for all taxa and the fact that there were few PAs greater than 20% of the species layer grid cell size (**Figure A5**), we present results/figures of the >20% PT; while sensitivity analyses relating to the choice of threshold are presented in the Supplementary Information (**Figure A4 & Table A1**). We also present the results of the >0% PT alongside to show a ‘best case’ scenario, which assumes any grid cell where PAs are present are effectively protecting evolutionary history, regardless of the percentage of the PA polygon that overlaps with the cell. In general, the locations of priority grid cells remained qualitatively similar when the analyses were run at each PT (**Figure A4**).

### 2.3. Phylogenetic analysis

The PD of each taxonomic group was calculated as the mean total length of all phylogenetic branches connecting all species present in the phylogeny, measured in billions of years (Gyr), across the sample of 25 phylogenetic trees. A phylogenetic branch was considered unprotected if no descendant species were deemed protected under our binary gap analysis. If at least one descendant species was considered protected, we considered all internal branches ancestral to that species to be protected. To determine the global distribution of unprotected PD, we summed the lengths of all branches in each unprotected grid cell that were unprotected globally (i.e. all grid cells in which all descendant species of the branch occur were unprotected).

To assess the similarity in the spatial distributions of PD between the different taxonomic groups we tested for correlations between the PD of each group across all grid cells when analyses were run under the >20% PT. The results for all other PT’s are provided in the supporting information (**Table A2**). We used a Moran’s I test (Gittleman & Kot 1990) to evaluate the data for spatial autocorrelation and ran pairwise correlations for all groups, corrected for spatial autocorrelation, using the R package ‘SpatialPack’ (Vallejos et al. 2018). A final pairwise correlation test was run between the spatial distributions of total amphibian PD and amphibian species richness. A Bonferroni correction for multiple testing was made to calculate the adjusted P-value at which to reject the null hypothesis.

To determine whether there was a significant difference in the proportion of unprotected PD within amphibian orders, we ran a one-way ANOVA. We then ran a Games-Howell post hoc analysis to identify pair-wise differences. All analyses were run in R version 3.5.3 (R Core Team 2019).

### 2.4. Priority grid cells

To investigate how increased coverage of the global PA network can capture unprotected PD, we identified the PD contribution of grid cells with the greatest unprotected amphibian PD. We identified the top 1% of grid cells with the highest levels of unprotected amphibian PD and designated them as ‘protected’. We then re-ran the gap analysis with the updated set of ‘protected’ grid cells and re-calculated the total amount of unprotected PD, identifying the gain in PD contribution of the newly protected percent of grid cells. We repeated this process until 50% of unprotected grid cells had been captured. For comparison, we repeated this analysis, this time selecting 1% of unprotected grid cells at random to be ‘protected’. As the complementary approach performed better than random for amphibians, we applied it to all taxonomic groups for comparison in the analyses.

Since the rate at which PD is captured declined markedly beyond 5% of grid cells (**Figure A6**), and in alignment with the methods of previous studies (Safi et al. 2013), we identified the top 5% of grid cells containing the largest amount of unprotected PD for each taxonomic group and highlighted them as ‘priority areas’. Grid cells that contain a large amount of unprotected PD for all taxonomic groups represent a good opportunity for maximum protection of terrestrial vertebrate PD by declaration of new PAs. The overlap between priority areas for each taxonomic group was quantified as the percentage of priority grid cells shared across all groups. Finally, to address the problem of feasibility of establishing new PAs, we identified the number of amphibian priority grid cells that are protected by lower IUCN management categories (III-VI), and which would benefit conservation of amphibian PD if it were possible to upgrade these PAs to IUCN management categories I or II.

## 3. Results

### 3.1. Global protection of amphibian PD

There were 212 379 PAs included in our analysis, including 19 637 (9.25%) PAs in IUCN management category I or II. Amphibians occurred in 11 482 grid cells (76.79% of global terrestrial land area). The percentage of grid cells meeting the PT declines rapidly as more stringent PTs are applied (**Table 1**), reflecting the fact that there are few PAs greater than 20% of the species layer grid cell size (100 × 100km; **Figure A7**). For all PT scenarios, more amphibian species were unprotected than either birds or mammals (**Figure 1A, Table A1**). Within amphibians, Caudata have the highest proportion of unprotected species (**Figure 1D**).

**Table 1:** Percentage of amphibian grid cells, terrestrial vertebrate species and phylogenetic diversity in IUCN PA management category I or II under different protection thresholds (PTs). Results for all other PTs are given in Table A1.

| Item description | Protection threshold (PT)<sup>a</sup> |  |
| --- | --- | --- |
|  | >0% | >20% |
| Amphibian grid cells protected (as a percent of total amphibian grid cells) | 6748 (58.77%) | 1192 (10.38%), see also Figure A7 |
| Amphibian species unprotected (as a percent of total species) | 1190 (20%) | 4579 (78%) |
| Caudata species | 92 (18%) | 489 (93%) |
| Anura species | 1063 (20%) | 3992 (77%) |
| Gymnophiona species | 26 (17%) | 119 (79%) |
| Bird species unprotected (as a percent of total species) | 650 (9%) | 2818 (39%) |
| Mammal species unprotected (as a percent of total species) | 478 (11%) | 2138 (49%) |
| Squamate species unprotected (as a percent of total species) | 1786 (19%) | 6581 (71%) |
| Amphibian PD<sup>b</sup> in billions of years unprotected (as a percent of total PD) | 14.56 (12%) | 75.88 (64%) |
| Caudata PD | 1.17 (12%) | 7.82 (78%) |
| Anura PD | 12.45 (12%) | 64.33 (63%) |
| Gymnophiona PD | 0.76 (12%) | 3.96 (64%) |
| Bird PD in billions of years unprotected (as a percent of total PD) | 3 (4%) | 16.44 (24%) |
| Mammal PD in billions of years unprotected (as a percent of total PD) | 2.7 (6%) | 14.68 (34%) |
| Squamate PD in billions of years unprotected (as a percent of total PD) | 15.22 (12%) | 72.24 (59%) |

<sup>a</sup>PT = Protection threshold
<sup>b</sup>PD = phylogenetic diversity

**Figure 1.**
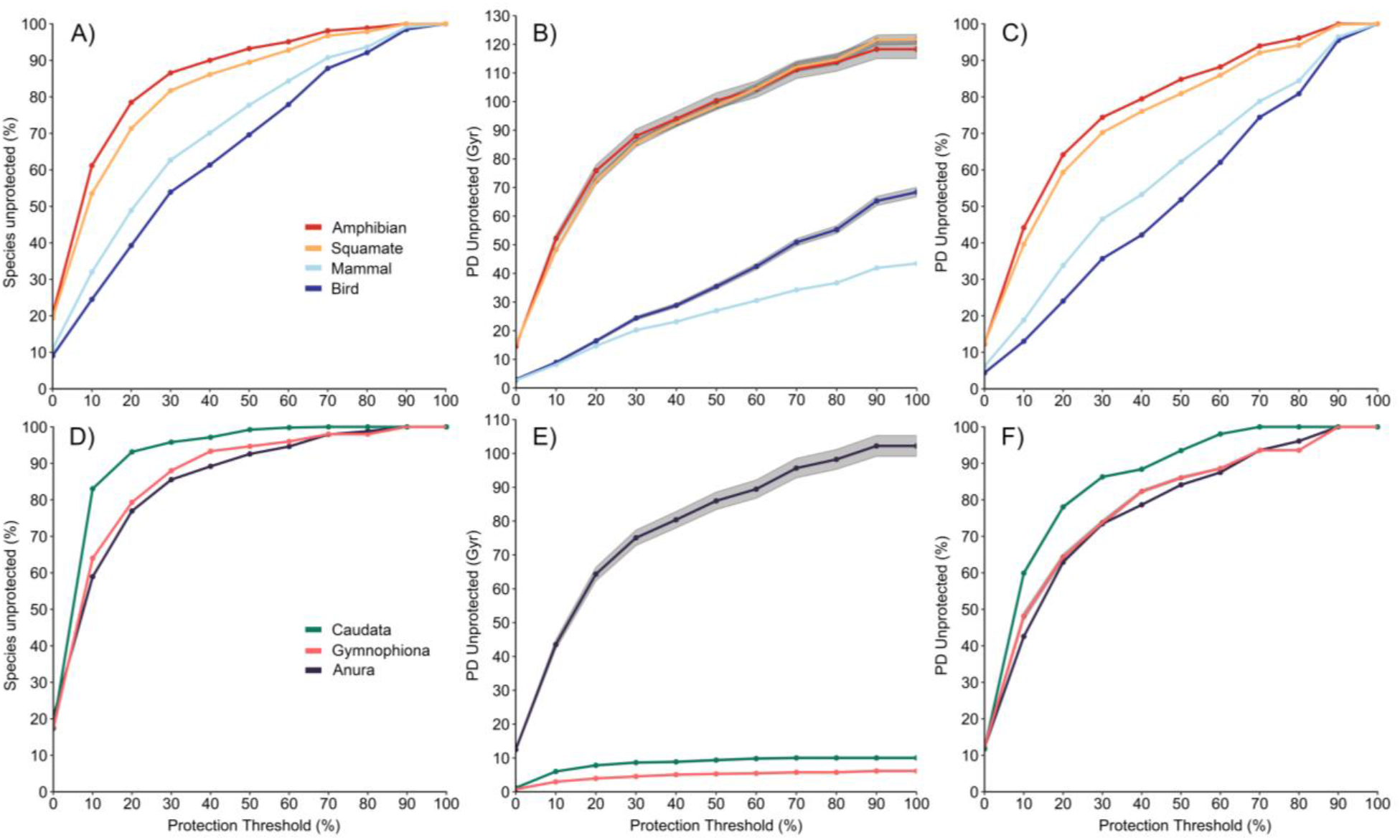
The percentage of species unprotected (**A, D**), the amount of unprotected PD in billions of years (Gyr; **B, E**) and the percentage of unprotected Phylogenetic Diversity (PD; **C, F**) for each taxonomic group (amphibians, squamates, mammals and birds; **A-C**) and amphibian Order (Anura, Caudata and Gymnophiona; **D-F**). Plots B, C, E and F show the mean +95% confidence interval (shaded grey) for the results of the 25 different phylogenetic trees, though the CI on plots C and F are narrow and barely visible behind the mean plot line.

Amphibians, followed closely by squamates, have the greatest amount of unprotected PD (**Figure 1B & 1C, Table 1**). The greatest difference in protection across taxa occurs at the lower-intermediate protection thresholds (20-50%; **Figure 1B & 1C**). The pattern was the same when all PA categories were used (Amount of unprotected PD in billions of years [as a percent of total PD]: Amphibian 41.33 [34.94%], Squamate 38.61 [31.70%], Bird 7.09 [10.36%], Mammal 6.42 [14.76%]) and areas identified as a priority remained largely the same (**Figure A8**).

Caudata has the greatest amount of unprotected PD of all amphibian orders (**Figure 1E & 1F, Table 1**) and the proportion of unprotected PD in Caudata is significantly higher than the proportion of unprotected PD in Anura and Gymnophiona (One-way ANOVA; F = 2446.3, p-value < 0.001, Games-Howell; p-value < 0.001). The results for all PTs are provided in the supporting information (**Table A3**). When all PA categories were used, Gymnophiona had the greatest proportion of unprotected PD, followed by Caudata (Amount of unprotected PD in billions of years [as a percent of total PD]: Gymnophiona 2.89 [47%], Caudata 4.17 [42%], Anura 34.19 [33%]) and areas identified as a priority for unprotected PD for all amphibian orders remained largely the same (**Figure A9**).

### 3.2. PD Distribution and Priority Areas

Global amphibian PD reflects amphibian richness patterns (Pearson’s correlation; r = 0.97, p-value < 0.001, **Figure A10**). Amphibian PD is poorly protected in the eastern United States, Central America, the Caribbean, the northern Andes and the Atlantic forests of Brazil (**Figure 2**). High levels of unprotected PD were also observed across Europe, Cameroon, Tanzania and South Africa, and Madagascar. In Asia, high levels of unprotected PD occur in the Western Ghats, southern China, Japan, Vietnam, Malaysian Borneo and the Philippines.

**Figure 2.**
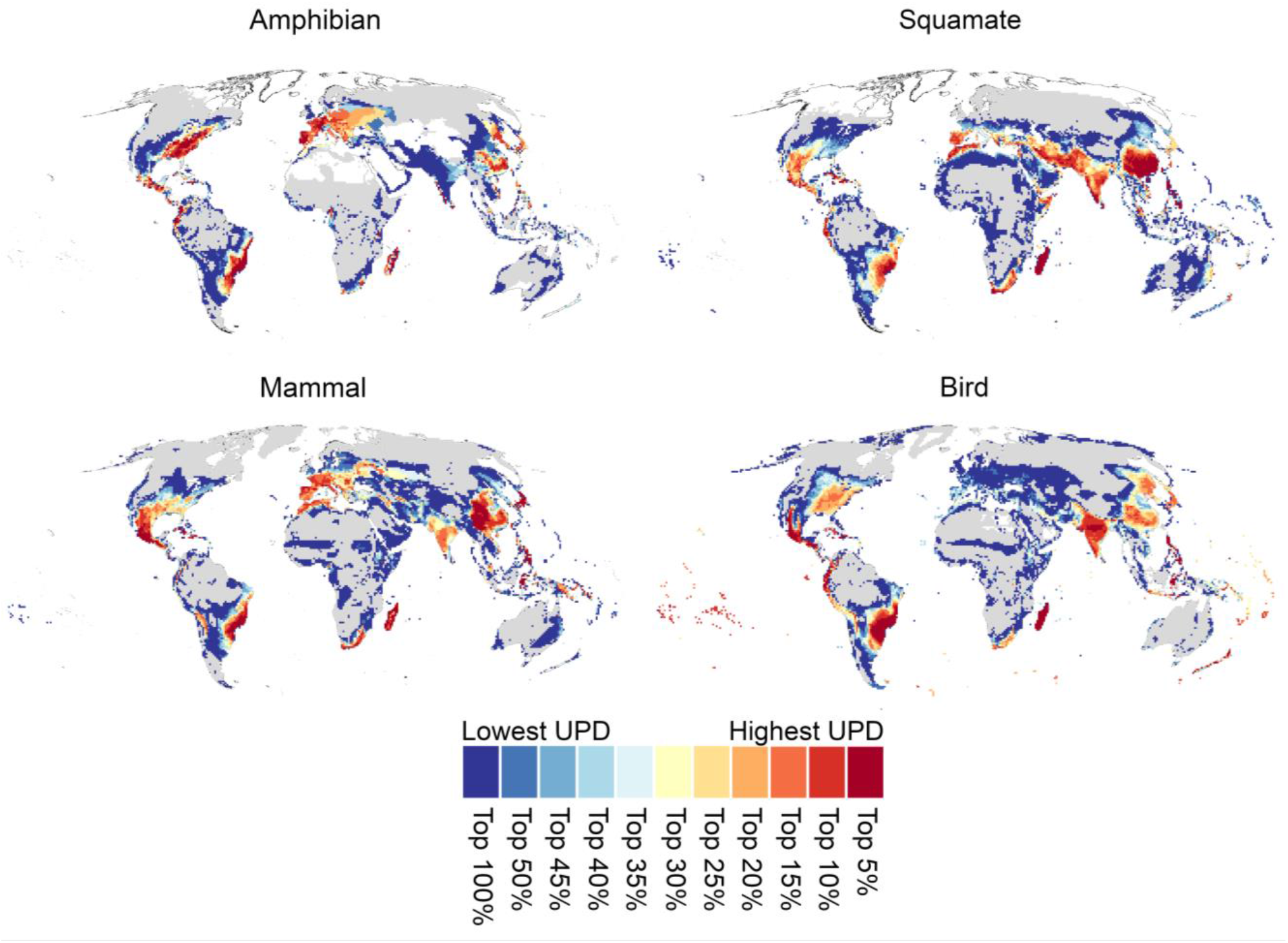
The distribution of unprotected phylogenetic diversity (UPD) for amphibians, mammals, birds and squamates, contrasted in ascending order of their level of protection. Dark grey area indicates the ranges of species that are protected in at least one grid cell. All maps were made under the >20% PT and results are the average across 25 separate phylogenetic trees. The maps for all other PT’s for amphibians are presented in the supporting information (**Figure A4**).

Many areas with high levels of unprotected amphibian PD harbour high levels of unprotected PD from all tetrapod groups, particularly across Central America, the Caribbean, the Atlantic forests of Brazil, Madagascar, the Western Ghats, and the Philippines. Overall, the spatial distribution of unprotected PD was strongly correlated across taxonomic groups (Pearson’s correlation; amphibian and squamate: r = 0.45, amphibian and mammal: r = 0.56, amphibian and bird: r = 0.57; all p-values < 0.001). However, correlations between non-amphibian groups were higher (Pearson’s correlation; birds and squamates: r = 0.63, birds and mammals: r = 0.65, mammals and squamates: r = 0.68; all p-value < 0.001; correlations for all PTs: **Table A2**).

The top 5% of grid cells of unprotected PD for amphibians, identified in our complementarity analysis (**Figure 2**), cover 289 grid cells (approximately 2 890 000 km2; 2.52% of all grid cells found to contain amphibians) and contained 52.73% of global unprotected amphibian PD (40.03 Gyr), and more than 29.43% of total amphibian PD (136 Gyr, Jetz & Pyron 2018). In addition, 67 (23% of 289) of these priority grid cells were found to be protected by lower category PAs (IUCN management categories III-VI), for which existing restrictions could potentially be increased. Therefore, establishing new PAs would be required in the remaining 222 priority grid cells (1.93% of all grid cells found to contain amphibians and 1.48% of all terrestrial grid cells), equalling approximately 2 220 000 km^2^ in area (1.74% of global land surface area).

Amphibian, bird, mammal and squamate priority grid cells were found to overlap by 3.32% (**Figure 3A**), with overlapping grid cells located across Hispaniola, the Atlantic forests of Brazil, and Madagascar. 11.31% of all priority grid cells across all taxonomic groups were unique to amphibians only, located across the Americas, Cameroon, Tanzania, southern Europe, the Western Ghats of India, and Vietnam (**Figure 3A**).

**Figure 3.**
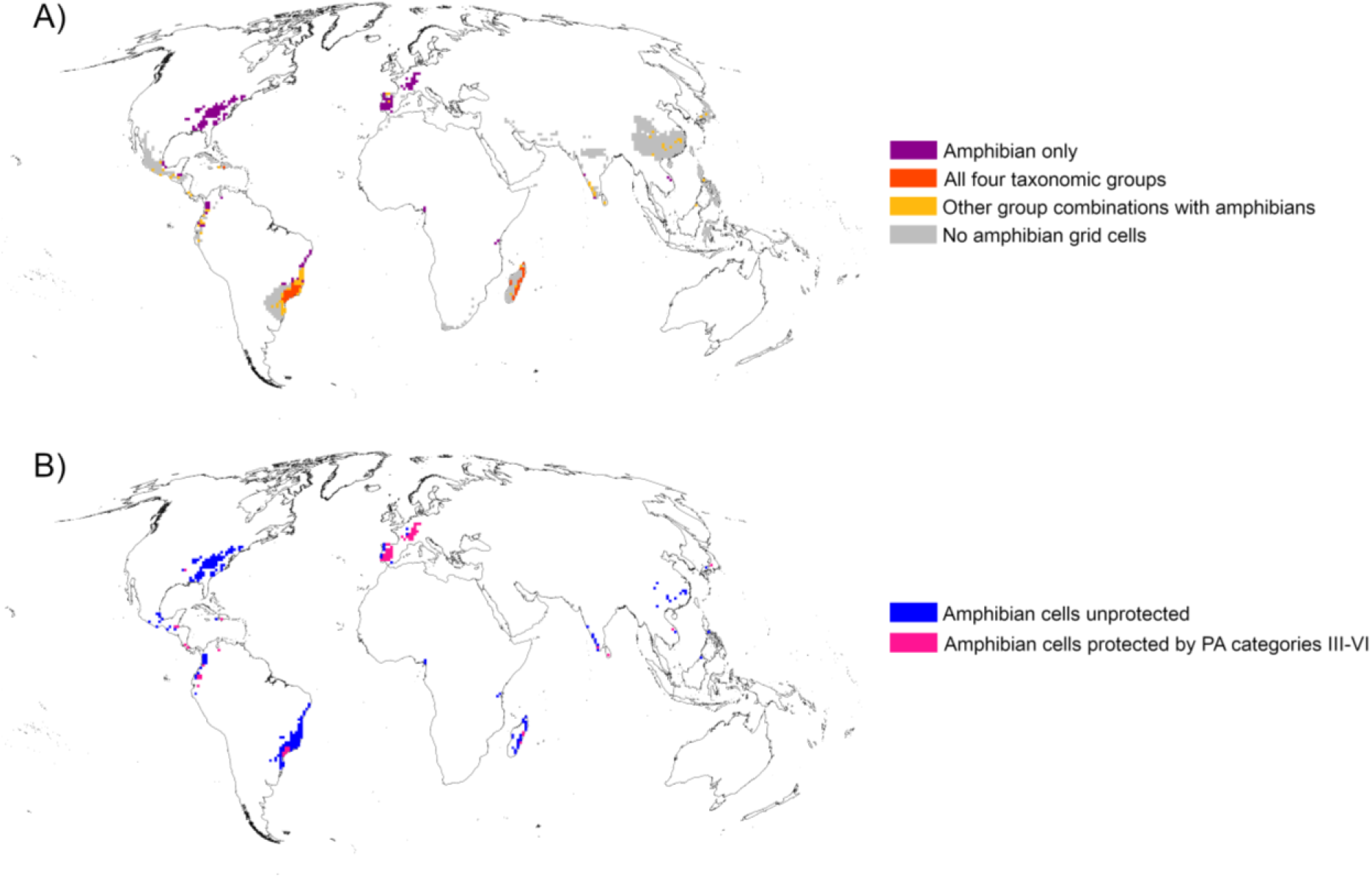
Priority grid cells when analyses were run at the >20% PT, calculated as the top 5% of grid cells containing the largest amount of unprotected phylogenetic diversity. **A)** The priority grid cells for all taxonomic groups. **B)** The Priority grid cells for amphibians only, with cells currently protected by PA categories III-VI highlighted.

There were spatial differences in the priority areas of unprotected PD for the different amphibian orders (**Figure 4**). For Caudata, priority areas occurred exclusively in the eastern United States. Priority areas of unprotected PD for Gymnophiona occurred in Colombia, Cameroon, Tanzania, the Seychelles and the Western Ghats of India. Priority areas for unprotected Anura PD were Central America, the northern Andes and the Atlantic forests of Brazil.

**Figure 4.**
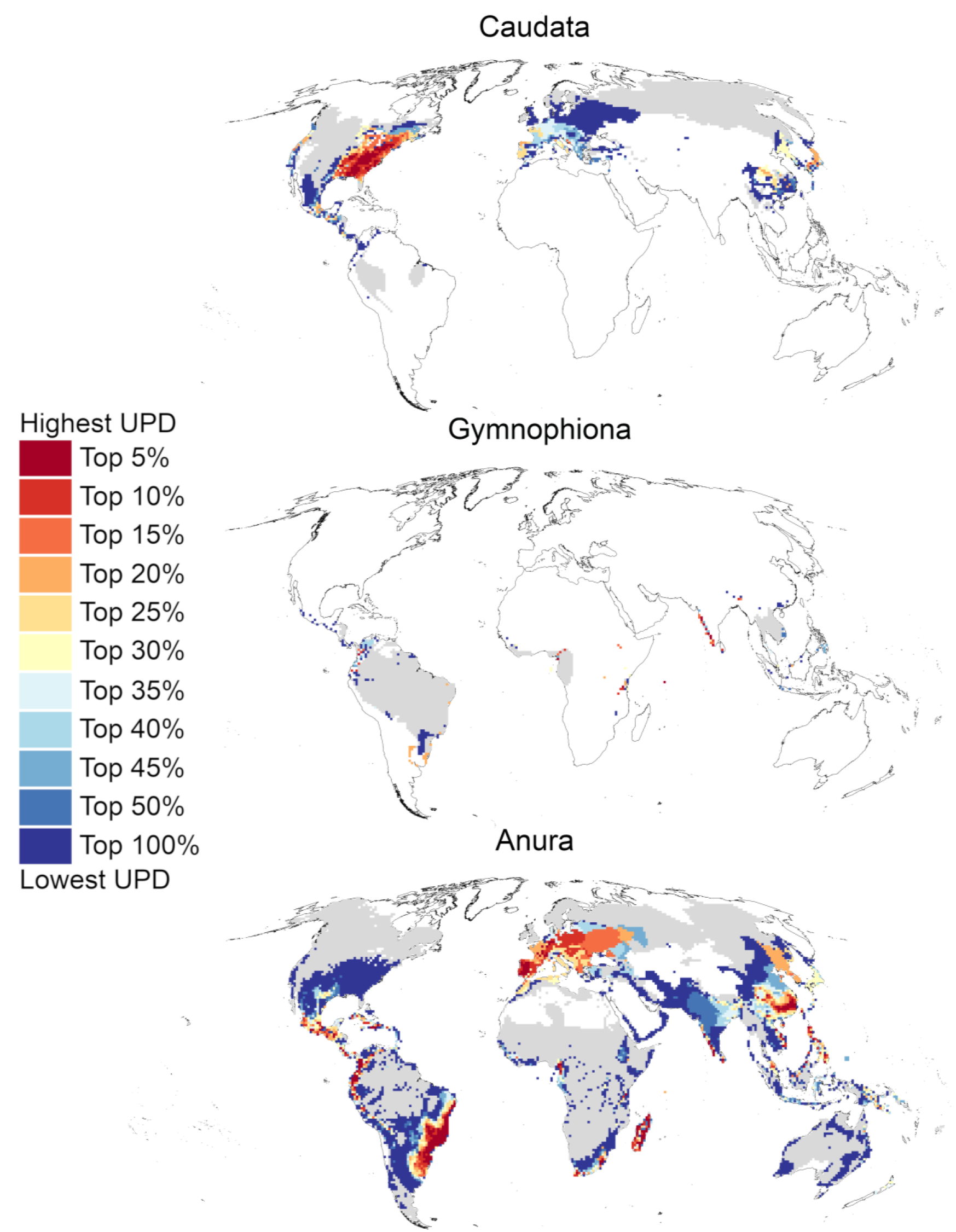
The distribution of unprotected phylogenetic diversity (UPD) for all three amphibian Orders, Anura, Gymnophiona and Caudata, contrasted in ascending order of their level of protection. Dark grey area indicates the ranges of species that are protected in at least one grid cell. All maps were made under the >20% PT and results are the average across 25 separate phylogenetic trees.

When the analyses were re-run using all PAs from all IUCN management categories (I-VI), 119 priority unprotected grid cells were identified for amphibians (**Figure A8**) and found to contain 17.66 Gyr of PD, the equivalent of 42.69% of unprotected PD (41.33 Gyr) and 12.99% of the PD of the entire clade. The spatial patterns of unprotected PD and the location of priority areas when all PA categories were included were consistent with the results when only PA management categories I and II were included (**Figure A8 & A9**). Results of analyses with 200 × 200 km resolution are consistent with those above and shown in the supporting information (**Figure A1 & A2**).

## 4. Discussion

Here we demonstrate that global amphibian PD is consistently under-protected relative to other terrestrial vertebrates. Despite being disproportionately threatened, conservation attention and action for amphibians and their associated habitat remains insufficient. Our gap analysis shows that, under our approach, 64% of all amphibian PD is not protected within the current terrestrial PA network. The lack of protection of the evolutionary history of Caudata and Gymnophiona is of particular conservation concern.

### 4.1. Global protection of amphibian PD

To reduce the omission and commission errors that occur when employing a binary setting of species protection, a broad interval range of PTs were tested. Under all PTs, from the most optimistic to the most conservative scenario (and at different spatial resolutions), amphibian PD was consistently the least well protected of all vertebrate groups and the relative levels of protection between the taxa remained the same across all thresholds, indicating that our results appear robust.

These findings reflect the well-established taxonomic bias in vertebrate conservation (Leader-Williams & Dublin 2000; Clark & May 2002; Rodrigues et al. 2004a, 2004b), and emphasises the need to prioritize these clades in future. Indeed, our analyses reflect recent findings that amphibians and squamates comprise significantly more PD than either mammals or birds and we stand to lose significantly more evolutionary history if they remain unprotected (Gumbs et al. 2020).

Among amphibians, Caudata have the largest proportion of evolutionary history at risk (78%), and all orders have a significant proportion of unprotected evolutionary history (>63%). When we ran our analyses to include all PA categories, Gymnophiona had the greatest proportion of unprotected PD, suggesting that Caudata PD is being better captured by PAs of lower management categories than that of Gymnophiona. Worryingly, previous research on data deficient (DD) amphibians has shown that the majority of their ranges (81%) lie completely outside of PAs (Nori & Loyola 2015), and a large proportion of amphibians, particularly Gymnophiona, are currently recognised as DD (Gymnophiona species: 55.7%, Anura: 20.3%, Caudata: 8.6%; IUCN 2020) or lack assessments entirely; future estimates of unprotected amphibian PD could therefore be even higher as more data becomes available.

### 4.2. PD Distribution and Priority Areas

Northern regions of South America contain the most amphibian species-rich area of the world (Stuart et al. 2004; Fritz & Rahbek 2012) and some parts appear to provide relatively strong protection for amphibian PD. However, priority regions for unprotected amphibian PD conservation occur in the northern Andes, the Atlantic forests of Brazil and the Eastern United States (US). Area-based conservation is complex and confounded by conflicting priorities; PAs tend to be designated in inaccessible places not wanted for other land uses and often suffer from a lack of international coordination (Visconti et al. 2019). In order to be effective, the post-2020 biodiversity framework must ensure spatial prioritisations to determine PAs value areas of high biodiversity importance and that various ecological and evolutionary processes are captured across borders (Visconti et al. 2019). The northern Andes and Atlantic forests are recognised as both Key Biodiversity Areas (KBAs) and UNESCO world heritage sites (Birdlife International 2020; UNESCO Institute for Statistics 2020). In combination with our high concentration of unprotected PD, it is clear that protecting these regions is of incredible importance.

At less stringent PTs, regions such as the US, the Northern Andes, Atlantic forests of Brazil, Madagascar and China remain priorities, whereas Europe, Japan and the Philippines appear to be more well protected. Previous studies into the protection of amphibian PD in Europe suggest that placement and habitat overlap of PAs with suitable habitat of amphibians is also important to be considered at a national and regional scale (Thuiller et al. 2015).

Priority grid cells for Caudata occurred exclusively in the US, particularly across the Appalachian mountain ranges, therefore extension of the US PA network can determine the future PD protection of the whole Caudata group. Gymnophiona PD, shown to occur in Cameroon, the Western Ghats of India and the Seychelles, provide the opportunity to capture a significant amount of PD for a lineage whose evolutionary history and ecology remains poorly understood (Stuart et al. 2004).

### 4.3. Taxonomic overlap of unprotected PD

Our analyses did highlight priority grid cells common to all clades; extending the PA network in the Atlantic forests of Brazil, Madagascar, and Hispaniola could capture large amounts of PD across all terrestrial vertebrate clades. For example, more than 13% of our priority grid cells for all terrestrial vertebrates were located in Madagascar, an island where PAs currently cover just 6% of its land (IUCN & UNEP-WCMC 2019). There was a strong correlation in unprotected PD patterns across the vertebrate groups, however, amphibian PD is not as strongly correlated with either mammal, bird or squamate PD, as they are with one another, highlighting the importance of identifying and conserving priority regions of unprotected amphibian PD in order to enhance overall terrestrial vertebrate PD protection. Priority grid cells of unprotected PD unique to amphibians predominantly coincide with areas already valued as Key Biodiversity Areas (Birdlife International 2020), re-emphasising the need for their protection.

### 4.4. Large PD gains possible for small PA increases

We have identified areas where increased protection could provide the largest gains in the conservation of amphibian evolutionary history. Increasing protection in only ~2.5% of the grid cells with amphibians could potentially capture ~30% (> 40 Gyr) of the evolutionary history of all amphibians and more than half of currently imperilled amphibian PD, offering an important opportunity to make large gains in amphibian PD conservation.

Our analysis has focused on PAs in IUCN management categories I and II because research has shown the reliable contribution of these PAs to biodiversity conservation (Jones et al. 2018). Amphibians are particularly susceptible to anthropogenic disturbance, therefore increasing restrictions in 23% (67/289 priority grid cells) of the priority regions found here, where there are PAs present but none that are managed as IUCN management categories I and II, will also benefit PD conservation. This is particularly relevant when considering the feasibility of transitioning to strict management from other classifications over establishing entirely new PAs and therefore presents an opportunity to also make rapid gains in PD conservation. We note that whether such a proportion of grid cells needs to be protected depends on the extent of suitable available habitat within the grid cells. Further work is needed to determine the opportunity cost of the priority regions and identify the optimal areas for protection to make sure that both cost efficiency and biodiversity gain are considered and weighted in the PA decision making (Carwardine et al. 2008a, 2008b; Venter et al. 2014).

### 4.5. Conclusion

PA expansion is high on the international agenda for the upcoming United Nations conferences as a tool to effectively protect biodiversity and the importance of conserving evolutionary history is gaining recognition. In order for this agenda to be adequately met there is an urgent need to prioritise under-represented species in conservation. Terrestrial vertebrates are an important group that require its own priorities that are better balanced across taxa. The assessment of terrestrial vertebrates in terms of their PD confirms that both amphibians and squamate reptiles are disproportionately under protected within the current global PA network. Relatively small increases in the PA network, as well as improvements of the existing network in key areas of high amphibian PD, could prevent a trajectory of excessive and unacceptable losses of evolutionary history and future options for humanity.

## 5. Acknowledgements

We thank the Department of Infectious Disease (DIDE) at Imperial College London for providing access to their high-performance computer to run these analyses. We also would like to thank Gerardo Martin and Trent Garner for their insightful comments. This work was supported by the Zoological Society London’s EDGE of Existence programme and the DIDE department at Imperial College London. R. Gumbs was funded by the NERC Science and Solutions for a Changing Planet Doctoral Training Programme (grant number NE/L002515/1).

## Notes

### Competing Interest Statement

The authors have declared no competing interest.

