## Supplementary Material for "Poor protection of amphibian evolutionary history reveals opportunities for global protected areas"

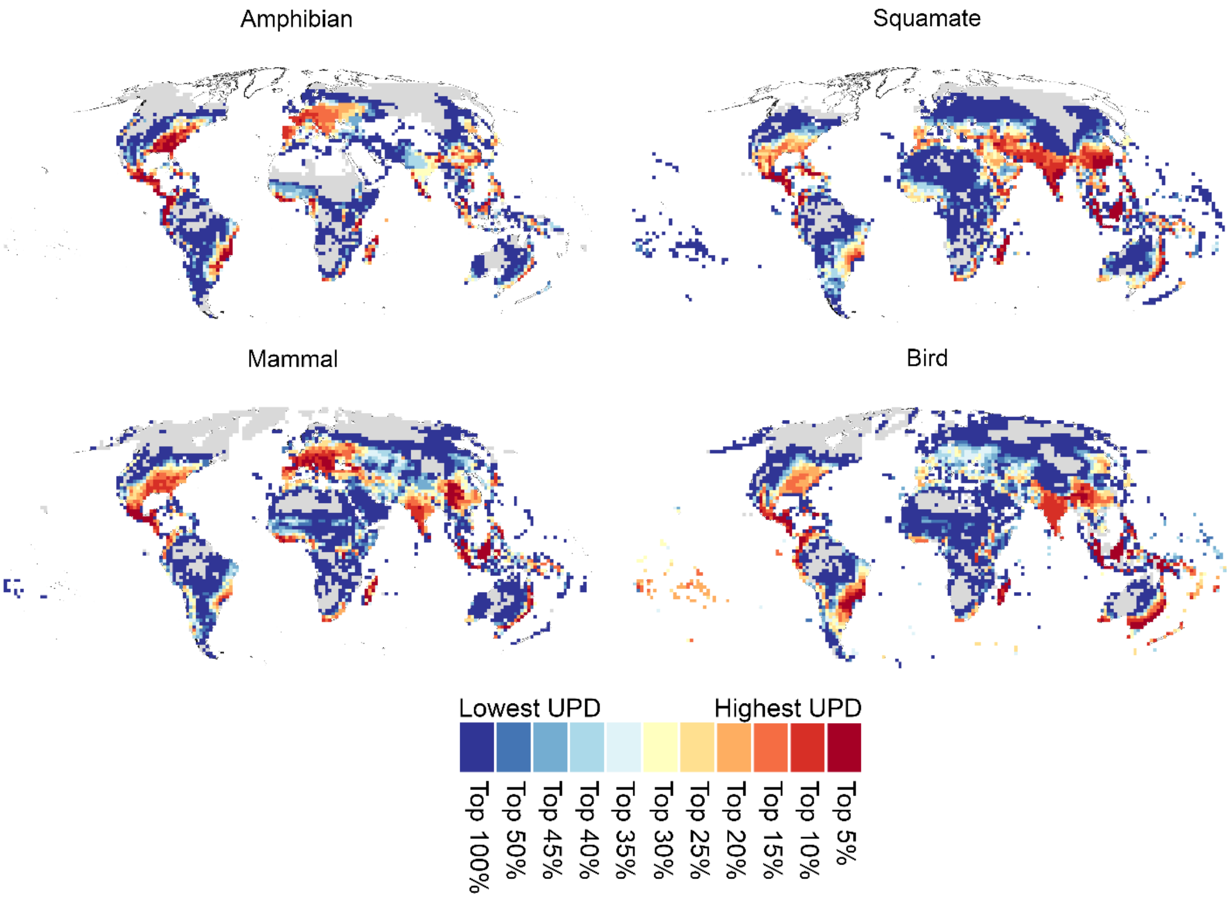


***Figure A1***. *The distribution of unprotected phylogenetic diversity (UPD) for amphibians, mammals, birds and squamates, when analyses were run at a 2 degree resolution and using protected areas (PAs) in IUCN management category I or II only, contrasted in ascending order of their level of protection. Dark grey area indicates the ranges of species that are protected in at least one grid cell. All maps were made under the >20% PT and results are the average across 25 separate phylogenetic trees.*

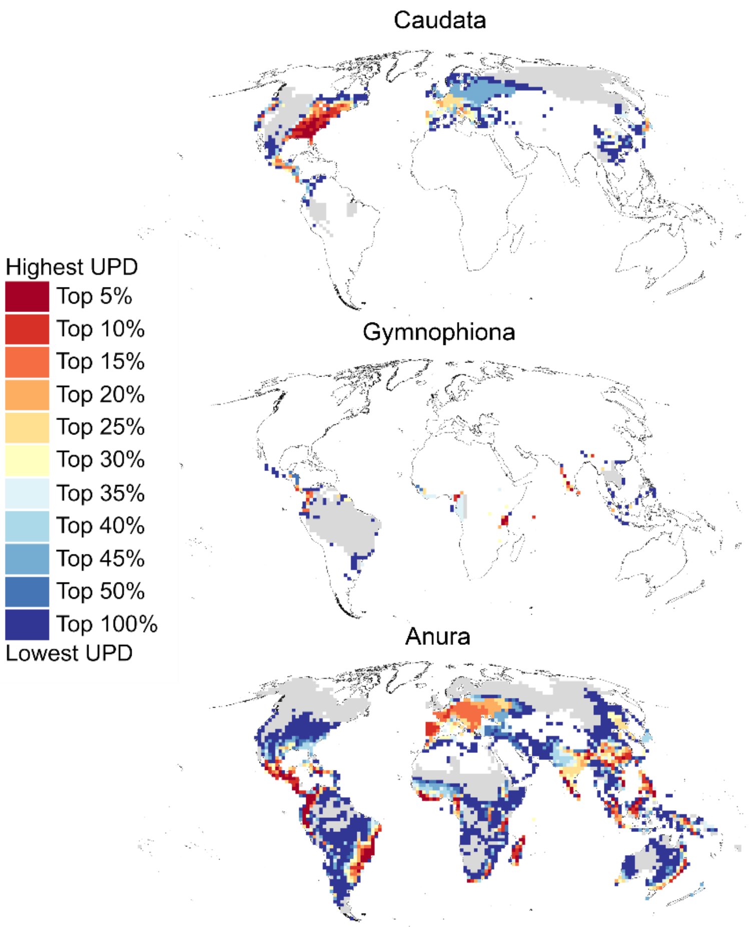


***Figure A2***. *The distribution of unprotected phylogenetic diversity (UPD) for all three amphibian Orders, Anura, Gymnophiona and Caudata, when analyses were run at a 2 degree resolution and using PAs in IUCN management category I or II only, contrasted in ascending order of their level of protection. Dark grey area indicates the ranges of species that are protected in at least one grid cell. All maps were made under the >20% PT and results are the average across 25 separate phylogenetic trees.*

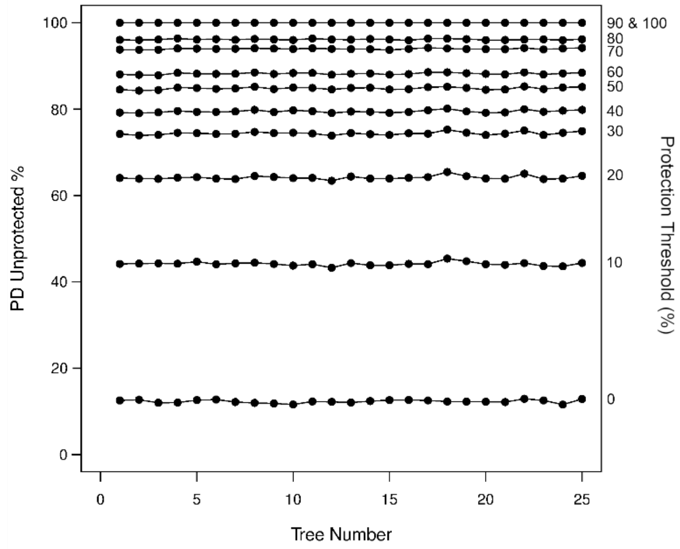

***Figure A3.*** *The proportion of unprotected phylogenetic diversity (PD) when amphibian analyses were run at a 1 degree resolution and using PAs in IUCN management category I or II only. Results are shown for each protection threshold (PT) and for each of the 25 separate phylogenetic trees.*

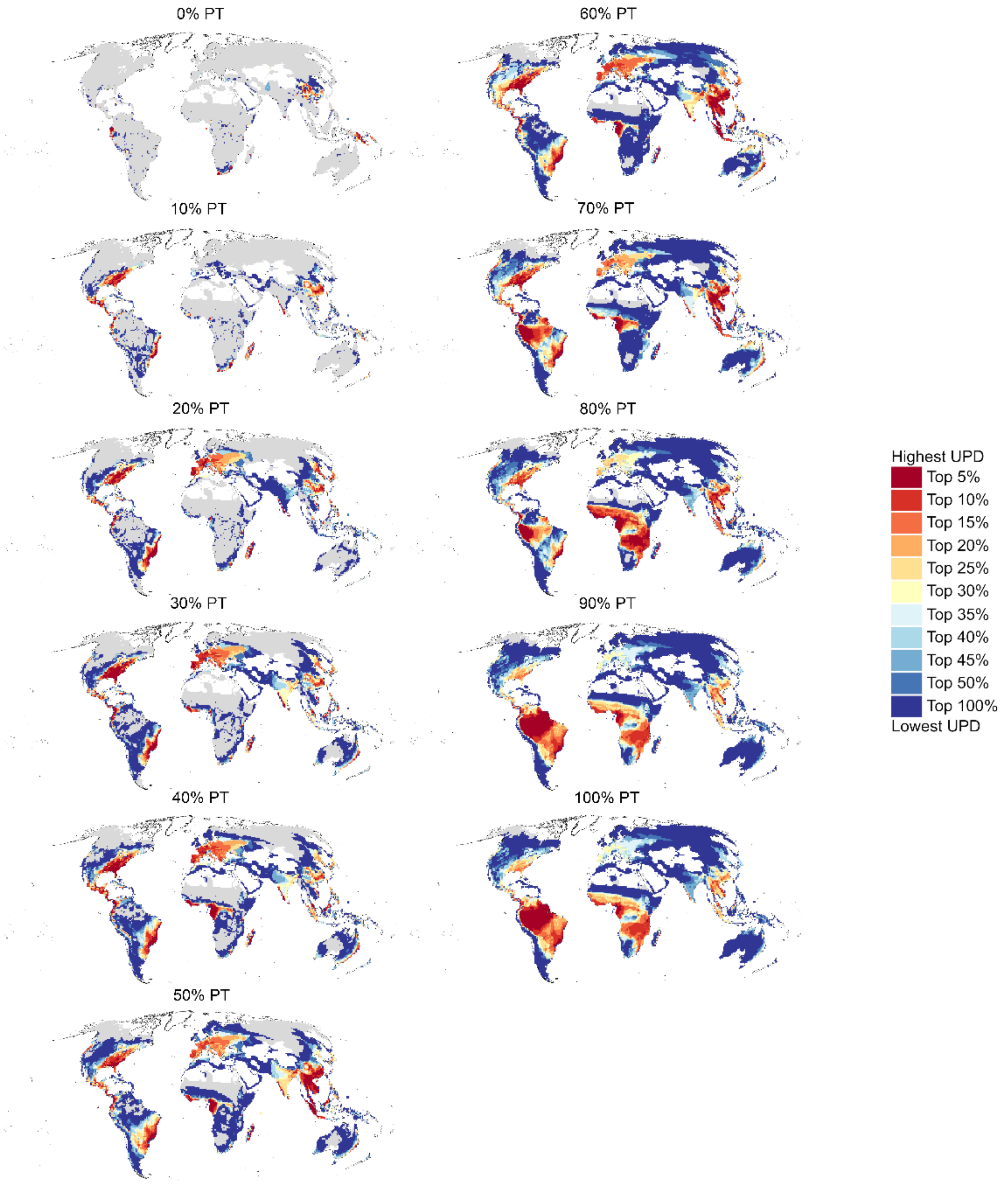

***Figure A4.*** *The distribution of unprotected phylogenetic diversity (UPD) for all amphibians when analyses were run using a 1 degree resolution and using PAs in IUCN management category I or II only, contrasted in ascending order of their protection threshold (PT). PT is defined by the proportion of the fine resolution raster with the protected area polygon that overlaps with the lower resolution species raster, e.g. 20% PT means that a grid cell is considered protected when at least 20% of the fine resolution raster polygon overlaps with the species raster. Dark grey area indicates the ranges of species that are protected in at least one grid cell. Results are the average across 25 separate phylogenetic trees.*

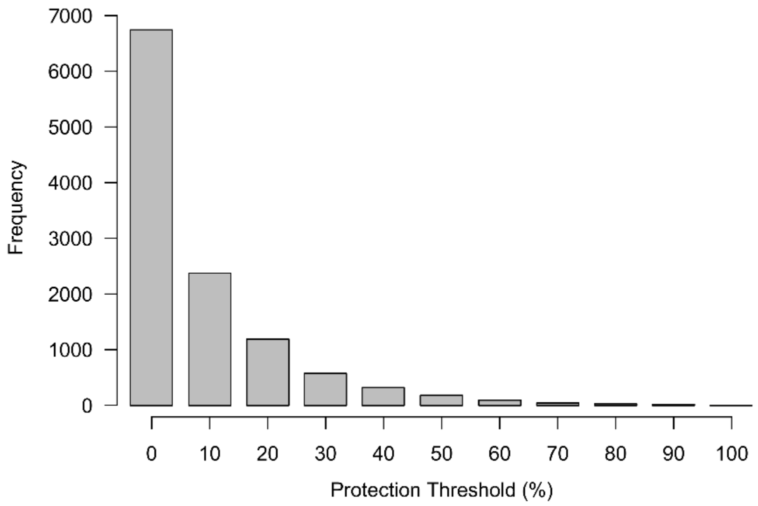


***Figure A5****. The frequency of grid cells that meet each protection threshold (PT), whereby PT refers to the percentage of overlap between the PA in IUCN management category I or II polygon and the grid cell.*

**Table A1:** The number of unprotected species and amount of unprotected phylogenetic diversity (PD) for each taxonomic group when analyses were run under each protection threshold, at a 1 degree resolution (~100 x 100km) and using PAs in IUCN management category I or II only. Results shown are the average (mean) results of 25 different phylogenetic trees.

| Taxa | Protection Threshold (%) | No. of spp. | Spp. Unprotected (%) | Total PD (Gyr) | PD unprotected (Gyr) | PD unprotected (%) |
|---|---|---|---|---|---|---|
| Bird | 0 | 7177 | 9.06 | 68.39 | 3.00 | 4.69 |
| | 10 | | 24.51 | | 8.91 | 13.45 |
| | 20 | | 39.26 | | 16.44 | 24.74 |
| | 30 | | 53.91 | | 24.40 | 36.23 |
| | 40 | | 61.33 | | 28.84 | 42.71 |
| | 50 | | 69.60 | | 35.44 | 52.65 |
| | 60 | | 77.89 | | 42.44 | 62.76 |
| | 70 | | 87.79 | | 50.88 | 74.84 |
| | 80 | | 92.11 | | 55.29 | 81.22 |

| | | | | | | |
|---|---|---|---|---|---|---|
| | 90 | | 98.45 | | 65.30 | 95.66 |
| | 100 | | 100.00 | | 68.39 | 100.00 |
| **Mammal** | 0 | 4374 | 10.93 | 43.46 | 2.70 | 6.20 |
| | 10 | | 31.98 | | 8.19 | 18.84 |
| | 20 | | 48.88 | | 14.68 | 33.78 |
| | 30 | | 62.67 | | 20.21 | 46.51 |
| | 40 | | 70.12 | | 23.15 | 53.25 |
| | 50 | | 77.73 | | 27.01 | 62.15 |
| | 60 | | 84.34 | | 30.52 | 70.22 |
| | 70 | | 90.72 | | 34.25 | 78.79 |
| | 80 | | 93.67 | | 36.68 | 84.40 |
| | 90 | | 99.06 | | 41.91 | 96.43 |
| | 100 | | 100.00 | | 43.46 | 100.00 |
| **Squamate** | 0 | 9229 | 19.35 | 121.76 | 15.22 | 12.50 |
| | 10 | | 53.49 | | 48.27 | 39.64 |
| | 20 | | 71.31 | | 72.24 | 59.32 |
| | 30 | | 81.71 | | 85.49 | 70.21 |
| | 40 | | 86.12 | | 92.55 | 76.00 |
| | 50 | | 89.47 | | 98.49 | 80.89 |
| | 60 | | 92.79 | | 104.62 | 85.92 |
| | 70 | | 96.73 | | 112.12 | 92.08 |
| | 80 | | 97.87 | | 114.64 | 94.15 |
| | 90 | | 99.99 | | 121.52 | 99.80 |
| | 100 | | 100.00 | | 121.76 | 100.00 |
| **Amphibian** | 0 | 5835 | 20.39 | 118.30 | 14.56 | 12.30 |
| | 10 | | 61.17 | | 52.23 | 44.15 |
| | 20 | | 78.47 | | 75.88 | 64.14 |
| | 30 | | 86.56 | | 87.98 | 74.38 |
| | 40 | | 90.01 | | 93.99 | 79.46 |
| | 50 | | 93.23 | | 100.31 | 84.80 |
| | 60 | | 95.10 | | 104.35 | 88.21 |
| | 70 | | 98.10 | | 111.14 | 93.95 |
| | 80 | | 98.89 | | 113.71 | 96.13 |
| | 90 | | 100.00 | | 118.30 | 100.00 |
| | 100 | | 100.00 | | 118.30 | 100.00 |
| **Anura** | 0 | 5188 | 20.49 | 102.21 | 12.45 | 12.18 |
| | 10 | | 58.94 | | 43.52 | 42.57 |
| | 20 | | 76.95 | | 64.33 | 62.92 |
| | 30 | | 85.54 | | 75.06 | 73.44 |
| | 40 | | 89.19 | | 80.39 | 78.65 |
| | 50 | | 92.58 | | 85.98 | 84.12 |
| | 60 | | 94.62 | | 89.45 | 87.52 |
| | 70 | | 97.92 | | 95.65 | 93.58 |
| | 80 | | 98.80 | | 98.23 | 96.11 |
| | 90 | | 100.00 | | 102.21 | 100.00 |

| | | | | | | |
|---|---|---|---|---|---|---|
| | 100 | | 100.00 | | 102.21 | 100.00 |
| | 0 | | 17.33 | | 0.76 | 12.29 |
| | 10 | | 64.00 | | 2.96 | 48.11 |
| | 20 | | 79.33 | | 3.96 | 64.30 |
| | 30 | | 88.00 | | 4.54 | 73.73 |
| | 40 | | 93.33 | | 5.07 | 82.34 |
| **Gymnophiona** | 50 | 150 | 94.67 | 6.15 | 5.30 | 86.06 |
| | 60 | | 96.00 | | 5.45 | 88.60 |
| | 70 | | 98.00 | | 5.76 | 93.60 |
| | 80 | | 98.00 | | 5.76 | 93.60 |
| | 90 | | 100.00 | | 6.15 | 100.00 |
| | 100 | | 100.00 | | 6.15 | 100.00 |
| | 0 | | 17.52 | | 1.17 | 11.73 |
| | 10 | | 83.05 | | 6.01 | 59.92 |
| | 20 | | 93.14 | | 7.82 | 78.05 |
| | 30 | | 95.81 | | 8.65 | 86.30 |
| | 40 | | 97.14 | | 8.86 | 88.39 |
| **Caudata** | 50 | 525 | 99.24 | 10.02 | 9.37 | 93.50 |
| | 60 | | 99.81 | | 9.83 | 98.05 |
| | 70 | | 100.00 | | 10.02 | 100.00 |
| | 80 | | 100.00 | | 10.02 | 100.00 |
| | 90 | | 100.00 | | 10.02 | 100.00 |
| | 100 | | 100.00 | | 10.02 | 100.00 |

**PD = phylogenetic diversity, Gyr = billions of years.*

**Table A2:** Results of the pairwise correlations between the spatial distributions of unprotected phylogenetic diversity (PD) values (billions of years) of each taxonomic group, when analyses were run under each protection threshold. A modified level of significance was accepted at 0.0125 (0.05 / 4; significance level / number of groups being tested)

| Pearson correlation test | Test results | Protection Threshold (%) | | | | | | | | | | |
|---|---|---|---|---|---|---|---|---|---|---|---|---|
| | | 0 | 10 | 20 | 30 | 40 | 50 | 60 | 70 | 80 | 90 | 100 |
| Amphibian & mammal | F-statistic | 30.41 | 69.78 | 202.50 | 185.45 | 303.21 | 110.67 | 123.84 | 95.93 | 71.76 | 75.86 | 94.75 |
| | df | 237.03 | 276.60 | 446.53 | 255.14 | 429.47 | 117.32 | 131.73 | 161.83 | 84.87 | 29.24 | 35.15 |
| | p-value | <0.001 | <0.001 | <0.001 | <0.001 | <0.001 | <0.001 | <0.001 | <0.001 | <0.001 | <0.001 | <0.001 |
| | r | 0.34 | 0.45 | 0.56 | 0.65 | 0.64 | 0.70 | 0.70 | 0.61 | 0.68 | 0.85 | 0.85 |
| Amphibian & bird | F-statistic | 26.19 | 47.12 | 343.28 | 248.39 | 296.43 | 103.52 | 125.76 | 48.35 | 81.93 | 81.93 | 97.50 |
| | df | 430.79 | 345.73 | 717.00 | 477.43 | 546.20 | 122.51 | 186.81 | 83.49 | 99.14 | 32.81 | 41.18 |
| | p-value | <0.001 | <0.001 | <0.001 | <0.001 | <0.001 | <0.001 | <0.001 | <0.001 | <0.001 | <0.001 | <0.001 |
| | r | 0.24 | 0.35 | 0.57 | 0.59 | 0.59 | 0.68 | 0.63 | 0.61 | 0.67 | 0.85 | 0.84 |
| Amphibian & squamate | F-statistic | 52.87 | 170.17 | 121.73 | 121.29 | 135.77 | 87.44 | 69.85 | 23.30 | 27.46 | 25.07 | 25.55 |
| | df | 444.69 | 437.16 | 483.71 | 376.89 | 510.97 | 138.32 | 129.75 | 85.30 | 85.71 | 31.58 | 31.31 |
| | p-value | <0.001 | <0.001 | <0.001 | <0.001 | <0.001 | <0.001 | <0.001 | <0.001 | <0.001 | <0.001 | <0.001 |
| | r | 0.33 | 0.53 | 0.45 | 0.49 | 0.46 | 0.62 | 0.59 | 0.46 | 0.49 | 0.67 | 0.67 |
| Mammal & bird | F-statistic | 13.33 | 106.54 | 631.46 | 337.50 | 769.35 | 278.14 | 286.11 | 190.40 | 214.14 | 201.18 | 218.51 |
| | df | 182.85 | 422.76 | 858.05 | 417.06 | 597.06 | 77.47 | 113.18 | 113.57 | 81.72 | 30.08 | 38.39 |
| | p-value | <0.001 | <0.001 | <0.001 | <0.001 | <0.001 | <0.001 | <0.001 | <0.001 | <0.001 | <0.001 | <0.001 |
| | r | 0.26 | 0.45 | 0.65 | 0.67 | 0.75 | 0.88 | 0.85 | 0.79 | 0.85 | 0.93 | 0.92 |
| Mammal & squamate | F-statistic | 26.59 | 92.03 | 358.86 | 155.64 | 333.74 | 302.33 | 184.07 | 188.80 | 119.14 | 79.94 | 49.05 |
| | df | 289.59 | 258.30 | 406.81 | 259.79 | 396.30 | 85.30 | 99.13 | 126.56 | 86.47 | 29.32 | 30.23 |
| | p-value | <0.001 | <0.001 | <0.001 | <0.001 | <0.001 | <0.001 | <0.001 | <0.001 | <0.001 | <0.001 | <0.001 |
| | r | 0.29 | 0.51 | 0.68 | 0.61 | 0.68 | 0.88 | 0.81 | 0.77 | 0.76 | 0.86 | 0.79 |
| Bird & squamate | F-statistic | 149.62 | 106.93 | 313.14 | 279.33 | 330.29 | 230.57 | 172.82 | 107.06 | 120.05 | 71.51 | 52.38 |
| | df | 671.29 | 477.74 | 465.16 | 373.24 | 305.83 | 75.36 | 90.82 | 55.49 | 72.52 | 30.35 | 37.93 |
| | p-value | <0.001 | <0.001 | <0.001 | <0.001 | <0.001 | <0.001 | <0.001 | <0.001 | <0.001 | <0.001 | <0.001 |
| | r | 0.43 | 0.43 | 0.63 | 0.65 | 0.72 | 0.87 | 0.81 | 0.81 | 0.79 | 0.84 | 0.76 |

**df = degrees of freedom. r = correlation coefficient.*

**Table A3:** Results of a One-way ANOVA test assuming unequal variances and the Games-Howell post-hoc analysis to identify pair-wise differences in the proportion (%) of unprotected phylogenetic diversity (PD) between the separate amphibian Orders, under each protection threshold. A modified level of significance was accepted at 0.01 (0.05 / 3; significance level / number of groups being tested).

| Protection Threshold (%) | Levene | | | One-way ANOVA | | | | Games-Howell Post hoc | | | |
|---|---|---|---|---|---|---|---|---|---|---|---|
| | df | F value | p-value | F value | num df | denom df | p-value | Order grouping | t | df | p-value |
| **0** | 2 | 11.21 | <.001 | 5.80 | 2 | 43.30 | 0.01 | Caudata-Anura | 3.31 | 45.10 | 0.005 |
| | | | | | | | | Gymno-Anura | 0.30 | 28.40 | 0.952 |
| | | | | | | | | Gymno-Caudata | 1.82 | 31.30 | 0.179 |
| **10** | 2 | 14.17 | <.001 | 2103.00 | 2 | 36.98 | <.001 | Caudata-Anura | 65.20 | 31.10 | <.001 |
| | | | | | | | | Gymno-Anura | 10.00 | 25.50 | <.001 |
| | | | | | | | | Gymno-Caudata | 19.80 | 33.60 | <.001 |
| **20** | 2 | 16.32 | <.001 | 2446.30 | 2 | 39.69 | <.001 | Caudata-Anura | 70.39 | 36.00 | <.001 |
| | | | | | | | | Gymno-Anura | 3.06 | 26.70 | 0.014 |
| | | | | | | | | Gymno-Caudata | 30.17 | 33.70 | <.001 |
| **30** | 2 | 21.58 | <.001 | 4041.30 | 2 | 41.10 | <.001 | Caudata-Anura | 90.11 | 40.20 | <.001 |
| | | | | | | | | Gymno-Anura | 0.81 | 26.40 | 0.701 |
| | | | | | | | | Gymno-Caudata | 35.23 | 30.10 | <.001 |
| **40** | 2 | 11.53 | <.001 | 2249.40 | 2 | 39.18 | <.001 | Caudata-Anura | 67.20 | 33.50 | <.001 |
| | | | | | | | | Gymno-Anura | 15.70 | 27.90 | <.001 |
| | | | | | | | | Gymno-Caudata | 25.40 | 40.30 | <.001 |
| **50** | 2 | 13.41 | <.001 | 2999.70 | 2 | 40.23 | <.001 | Caudata-Anura | 78.09 | 35.50 | <.001 |
| | | | | | | | | Gymno-Anura | 9.08 | 28.40 | <.001 |
| | | | | | | | | Gymno-Caudata | 36.99 | 39.40 | <.001 |
| **60** | 2 | 15.69 | <.001 | 11918.00 | 2 | 42.56 | <.001 | Caudata-Anura | 154.17 | 39.20 | <.001 |
| | | | | | | | | Gymno-Anura | 8.37 | 31.30 | <.001 |
| | | | | | | | | Gymno-Caudata | 91.47 | 41.60 | <.001 |
| **70** | 2 | 30.51 | <.001 | NA | 2 | NA | NA | Caudata-Anura | 395.10 | 24.00 | <.001 |
| | | | | | | | | Gymno-Anura | 0.26 | 31.80 | 0.964 |
| | | | | | | | | Gymno-Caudata | 160.95 | 24.00 | <.001 |
| **80** | 2 | 31.49 | <.001 | NA | 2 | NA | NA | Caudata-Anura | 252.40 | 24.00 | <.001 |
| | | | | | | | | Gymno-Anura | 32.20 | 35.10 | <.001 |
| | | | | | | | | Gymno-Caudata | 160.90 | 24.00 | <.001 |
| **90** | 2 | NA | NA | NA | 2 | NA | NA | Caudata-Anura | NA | NA | NA |
| | | | | | | | | Gymno-Anura | NA | NA | NA |
| | | | | | | | | Gymno-Caudata | NA | NA | NA |
| **100** | 2 | NA | NA | NA | 2 | NA | NA | Caudata-Anura | NA | NA | NA |
| | | | | | | | | Gymno-Anura | NA | NA | NA |
| | | | | | | | | Gymno-Caudata | NA | NA | NA |

**df = degrees of freedom.*

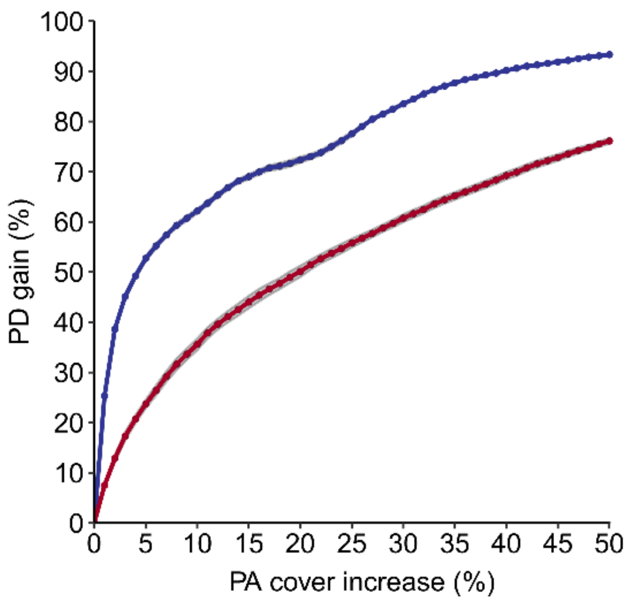


***Figure A6.*** *The rate at which amphibian PD is captured (%) when 1-50% of unprotected grid cells are protected. The blue line shows when 1-50% of the highest scoring unprotected grid cells are protected. The red line shows when 1-50% of randomly selected grid cells are protected. Each line shows the mean +95% confidence interval (shaded grey) for the results of the 25 different phylogenetic trees.*

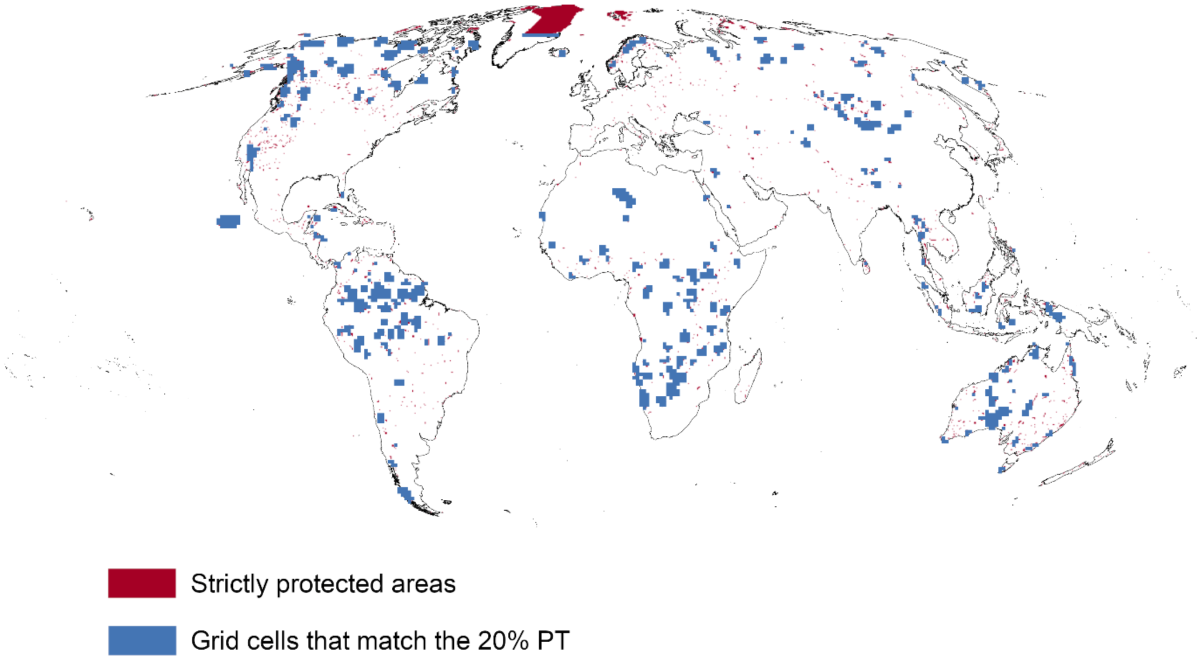


***Figure A7****. Global distribution of PAs in IUCN management category I or II when rasterized at a resolution of 2.5 x 2.5 degrees (red) and 100 x 100 km (blue). The blue raster has been subset to only include grid cells that match the >20% protection threshold (PT), meaning at least 20% of the grid cell overlaps with a PA in IUCN management category I or II.*

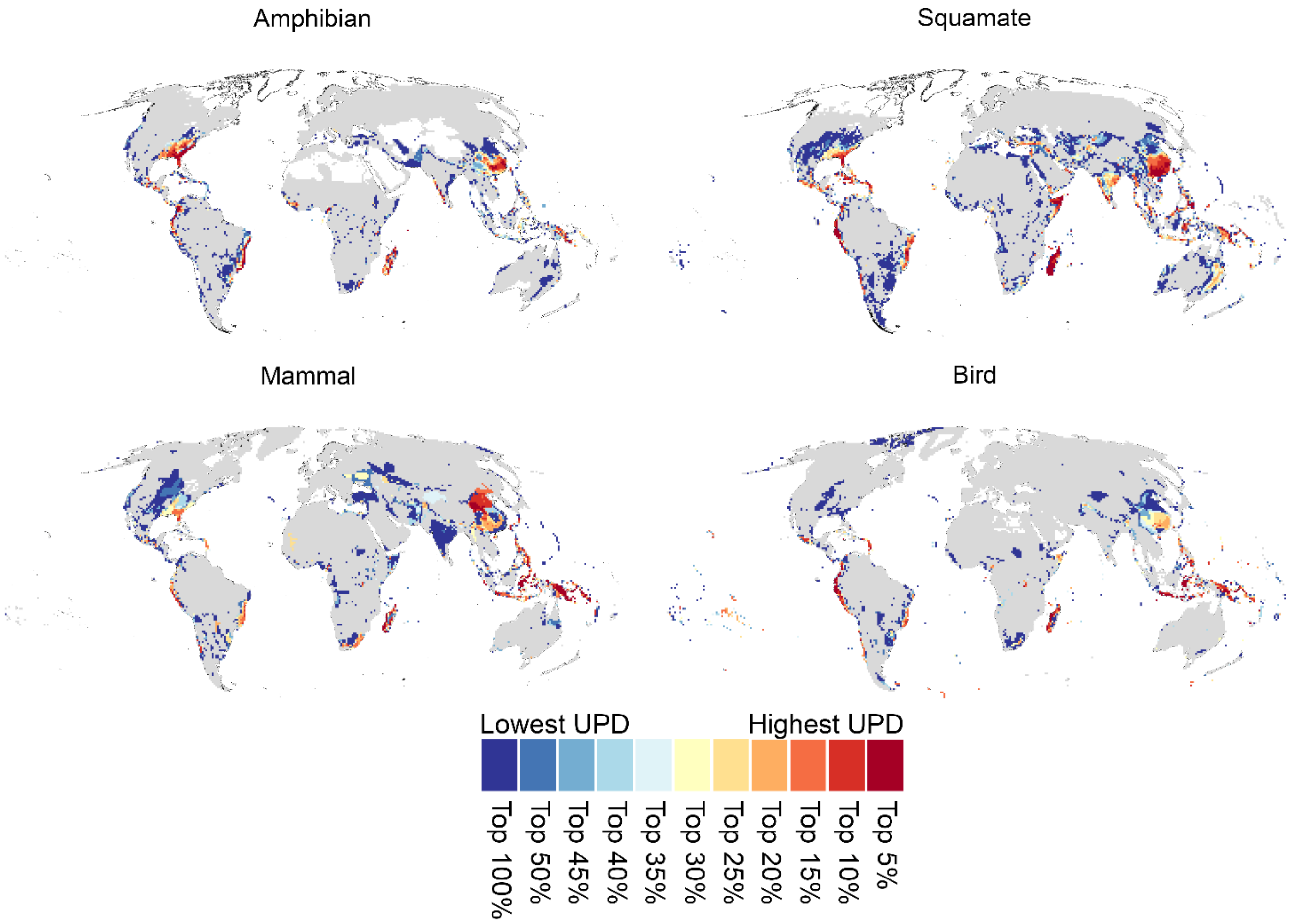


***Figure A8**. The distribution of unprotected phylogenetic diversity (UPD) for amphibians, mammals, birds and squamates, when analyses were run at a 1 degree resolution and using all IUCN categories of protected areas (including management categories I-VI and all PAs labelled as 'Not Applicable', 'Not Assigned' and 'Not reported'). Maps are contrasted in ascending order of their level of protection. Dark grey area indicates the ranges of species that are protected in at least one grid cell. All maps were made under the >20% PT and results are the average across 25 separate phylogenetic trees*.

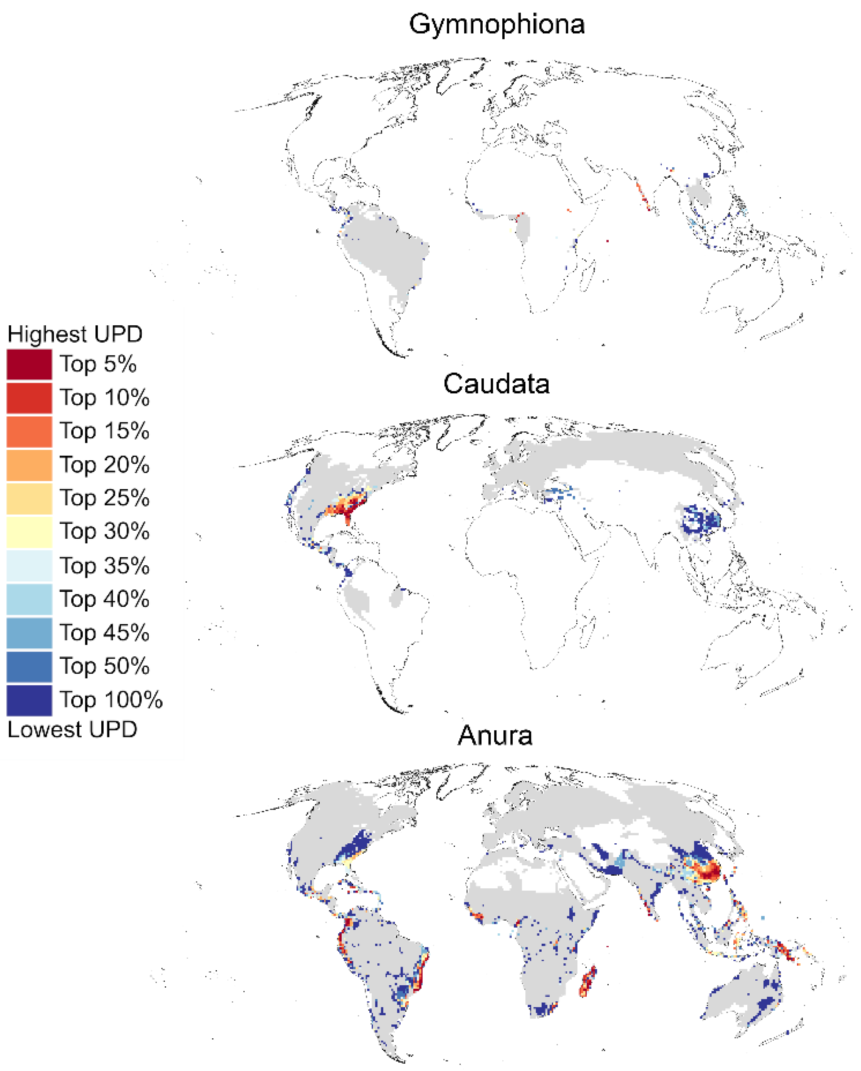


***Figure A9***. *The distribution of unprotected phylogenetic diversity (UPD) for all three amphibian Orders, Anura, Gymnophiona and Caudata, when analyses were run at a 1 degree resolution and using all IUCN categories of protected areas (including management categories I-VI and all PAs labelled as 'Not Applicable', 'Not Assigned' and 'Not reported'). Maps are contrasted in ascending order of their level of protection. Dark grey area indicates the ranges of species that are protected in at least one grid cell. All maps were made under the >20% PT and results are the average across 25 separate phylogenetic trees.*

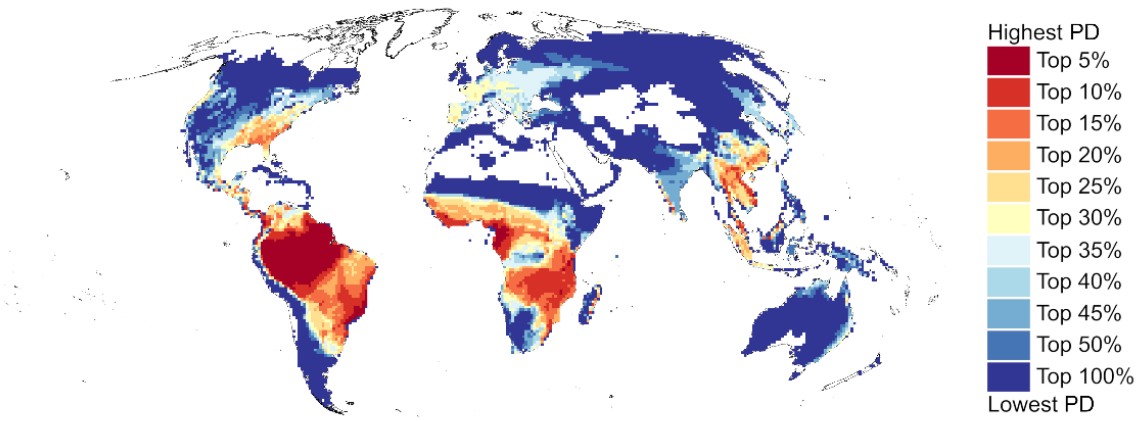


***Figure A10.*** *The global distribution of amphibian phylogenetic diversity (PD) when analyses were run using a 1 degree resolution. Results are the average across 25 separate phylogenetic trees.*
